# Differential Expression Gene Explorer (DrEdGE): A tool for generating interactive online data visualizations for exploration of quantitative transcript abundance datasets

**DOI:** 10.1101/618439

**Authors:** Sophia C. Tintori, Patrick Golden, Bob Goldstein

## Abstract

As the scientific community becomes increasingly interested in data sharing, there is a growing need for tools that facilitate the querying of public data. Mining of RNA-seq datasets, for example, has value to many biomedical researchers, yet is often effectively inaccessible to non-genomicist experts, even when the raw data are available. Here we present DrEdGE (dredge.bio.unc.edu), a free Web-based tool that facilitates data sharing between genomicists and their colleagues. The DrEdGE software guides genomicists through easily creating interactive online data visualizations, which colleagues can then explore and query according to their own conditions to discover genes, samples, or patterns of interest. We demonstrate DrEdGE’s features with three example websites we generated from publicly available datasets—human neuronal tissue, mouse embryonic tissue, and a *C. elegans* embryonic series. DrEdGE increases the utility of large genomics datasets by removing the technical obstacles that prevent interested parties from exploring the data independently.

## Background

Data sharing, in the interest of transparency, openness, and reproducibility, is increasingly becoming embraced by the scientific community^1–6^. The practice of data sharing allows colleagues to independently reproduce and verify analyses, and to build future research upon them more reliably. In burgeoning big data fields such as genomics, sharing enables data mining, wherein other researchers can perform their own analyses on large, multivalent datasets that may have only been used for a small fraction of possible applications at the time of initial publication^7–9^.

Currently, the primary method for sharing data is to publish raw data on databases, such as the NCBI Gene Expression Omnibus, the NCBI Sequence Read Archive, DNA Data Bank of Japan, European Nucleotide Archive, Figshare, and Dryad^10–15^. These raw data repositories are critical, as they allow for independent analyses from entirely unprocessed material. Also critical are tools for sharing partially processed data in an interactive format. Such tools facilitate communication with non-genomicist collaborators and colleagues by allowing them to explore the data without having to reprocess them starting from the rawest form, which can be considerably, often prohibitively, time consuming. Without such tools, the data may be available but are not effectively accessible to most non-genomicist researchers^16,17^.

Genome browsers (such as those of UCSC^18^, Ensembl^19^, and NCBI^20^) represent one such type of interactive tool that allows researchers to explore partially processed data. Users can browse sequencing read density at each nucleotide in the genome from a given experiment, alongside annotation of genomic features such as introns and exons, GC content, repeats, and conservation scores. While this kind of tool is critically useful for visually identifying patterns in the data, users cannot generate statistical results, nor easily ask biological questions that span more than one locus. As sequencing costs continue to drop^21^ and multiplexing technology becomes more sophisticated^22–24^, the rate at which large multivalent datasets are generated is increasing^25^. In turn, the need for tools that make these datasets accessible continues to grow.

To address this need, we have created the Differential Expression Gene Explorer (DrEdGE, dredge.bio.unc.edu), a web-based interactive data visualization tool. DrEdGE allows genomics researchers to share their datasets in a readily accessible, queryable format, and allows collaborators or colleague to explore the datasets and identify genes or samples of interest. DrEdGE’s flexible design can be used to visualize any number of sample types, and any type of quantitative unit (transcripts, DNA fragments, proteins, etc.), using any statistical model for differential expression. The user can interact with three different data representations on a DrEdGE website—a differential expression plot, a statistical table, and an experiment-wide heat map—which each feed into the other representations, creating an iterative workflow that can be cycled through repeatedly for the continuous fine tuning or elaboration of hypotheses. We provide a video to illustrate the features of a DrEdGE website (https://vimeo.com/336521735/c1e473b81e), and a video to guide researchers in creating their own DrEdGE website (https://vimeo.com/336521569/816b9585d8).

## Results

### GENERAL PROPERTIES

DrEdGE is designed to connect two groups of researchers: those who create, process, and publish genomic datasets, and those interested in asking biological questions of genomic data (who may or may not have specialized computational genomics skills). In this manuscript we distinguish between these two types of users. The first user—the author—makes an interactive DrEdGE website by importing their dataset. The author can publish this website alongside their data and analyses in a peer-reviewed journal, or he or she can share it privately with colleagues. The second user—the reader—explores the DrEdGE website, querying the data with his or her own biological questions.

We will first describe the DrEdGE Web application, as used by the reader for exploration and analysis of biological data. Then, we will describe a few sample datasets used to make sample DrEdGE websites. Finally, we will describe the process by which the author prepares and uploads a new dataset to create his or her own novel DrEdGE website.

### PRIMARY FEATURES OF DREDGE VISUALIZATION

The DrEdGE website displays gene expression data based on two variables: (1) units of genetic material (i.e. cDNA, genome fragments, proteins—because we showcase RNA-seq experiments in examples below, we will call these units "transcripts"), and (2) sets of experimental replicates (i.e. tissue types, chemical treatments, stages in a time course—we will call these units "treatments") within an experiment.

Three visual elements make up the visualization: (1) a dot plot displaying relative abundance of transcripts between two treatments, (2) a data table listing transcript abundance measurements and statistics, and (3) a heat map displaying the relative abundances of transcripts across every treatment in the experiment. Results from each data representation can be used as input for another representation, allowing the user to build upon their hypotheses in an iterative fashion. A video description of these features is available at https://vimeo.com/336521735/c1e473b81e.

#### MA plot

The reader begins an investigation by selecting two treatments to compare (Figure 1A). Treatments are selected from either a vector graphic representing the dataset, or a simple dropdown menu in which all treatments are listed. The treatments will then populate the rest of the visualization, starting with the MA (log ratio *M* by mean average *A*) plot, which describes differential abundance (in RPKM or otherwise normalized values) between the two treatments for all transcripts (Figure 1B). These plots are commonly used to visualize how abundant a transcript is and how specifically it is enriched in one treatment over another. Each dot on this plot represents one or more transcripts. Transcripts in the top half are more abundant in the treatment specified above the plot, and those on the bottom half are more abundant in the treatment specified below. Transcripts to the right of the plot have a greater average abundance than those to the left. The reader selects transcripts in an area of the MA plot by brushing (clicking and dragging) with the mouse, and transcripts selected in this way then populate further elements of the visualization. The reader can filter out transcripts by p-value of differential abundance by sliding the p-value cutoff selector (Figure 1C).

**Figure 1:**
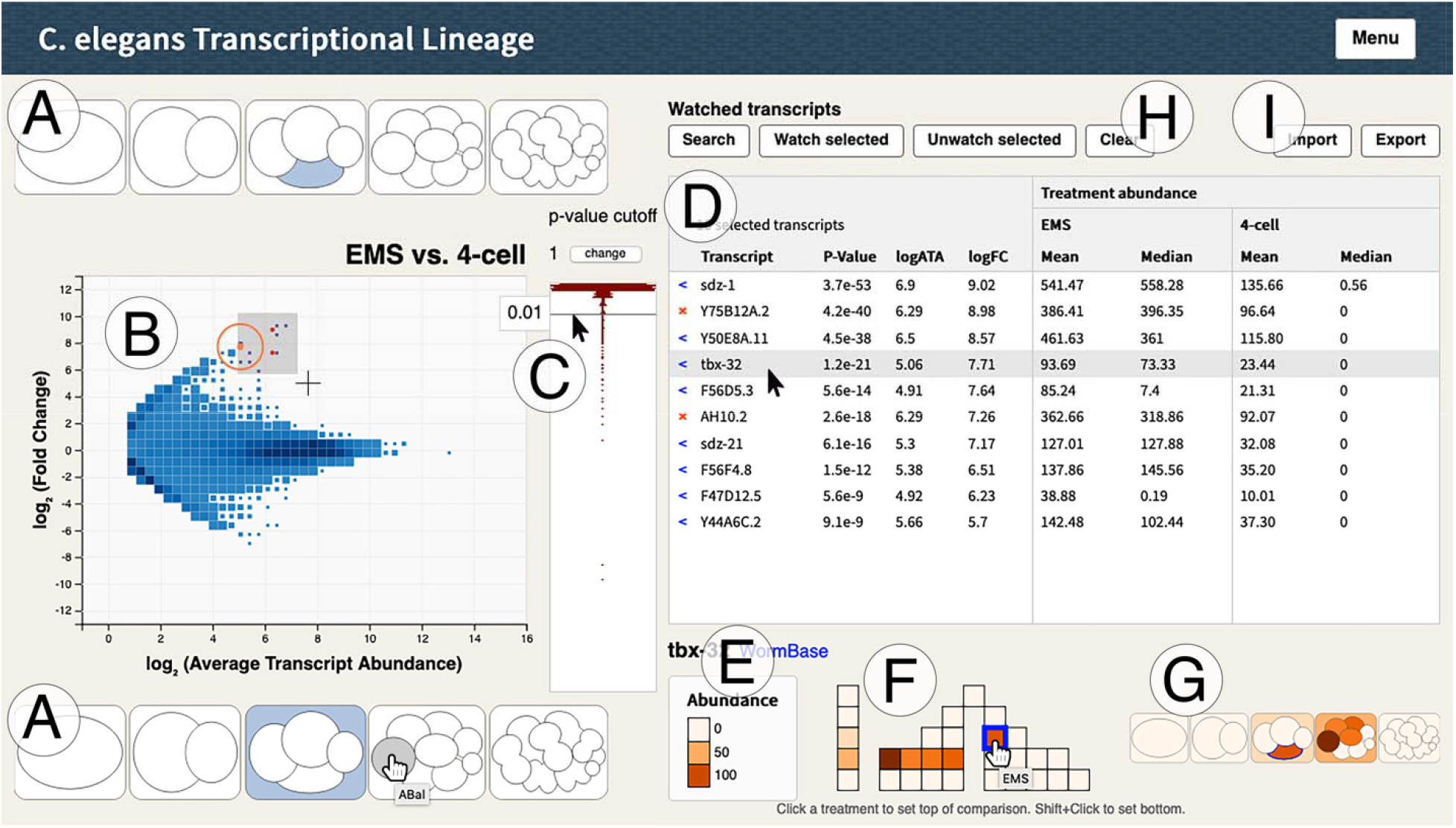
Layout of DrEdGE visualization. A) The user can select two treatments to compare, using icons or a drop column. B) An MA plot shows differential expression between two treatments. C) A slider that filters transcripts out of the MA plot by p-value. D) A table shows statistical values of transcripts highlighted from the MA plot. E) A heat map shows relative transcript abundance across all treatments for a transcript selected from the table, graphically represented as F) a grid of square icons, G) an illustration, or both. H) A curated list of transcripts can be saved, added to, pruned, or cleared, through multiple analyses. I) Lists of transcripts can be imported or exported to save, share with colleagues, or interface with other methods and analyses in the user’s analytical pipeline. Data is from Tintori et al, 2016^27^.

#### Data table

Transcripts highlighted on the MA plot appear in the table to the right (Figure 1D). This table displays the fold change values between the two treatments selected, p-values for fold change, and transcript abundance values for each treatment. The reader can then sort the transcripts by name, average abundance, ratio of abundances between the two treatments, or p-value. Abundance levels for each treatment are available both as mean and median, which allows some insight into the distribution of data across replicates.

The reader can curate a list of transcripts of note by adding them to a “watched” list in the data table. These watched transcripts will persist in the table, highlighted by red dots, when no area is brushed and when new treatments are selected for comparison. Watched transcripts can be added or removed, allowing the reader to curate a list of transcripts that fit multiple criteria spanning several pairwise comparisons. To add transcripts one-by-one to the watched list, the reader can either click the arrow next to a selected transcript name on the table, or type transcripts in a search bar. To add groups of transcripts to the watched list, the user can upload a comma delineated file with one transcript name on each line, or the user can click “watch selected,” which will add all transcripts currently brushed over on the plot. To remove transcripts the reader either clicks the X next to each transcript on the table, brushes over a region of the MA plot then clicks “remove selected,” or clicks “remove all.” At any point the reader can export their current list of watched transcripts.

As the user hovers over or clicks transcripts in the data table, the visualization updates to display additional information. First, an orange circle appears in the MA plot around that transcript’s position. Second, a heat map diagram is populated under the table, showing the relative abundance of the transcript for all treatments within the dataset.

#### Heat map

The heat map (Figure 1E) graphically represents abundances of a single transcript across all experimental treatments. This feature allows a reader to examine the experiment-wide context for a transcript that was selected based on just a pairwise comparison. If the heat map reveals that another treatment besides the two selected for the MA plot is of interest, the reader can click on that treatment’s icon to replace the top or bottom treatment in the MA plot. Heat maps may be shown as a grid of boxes (as in Figure 1F), as illustrated icons (as in Figure 1G), or both.

#### Iterative analysis

Data points from each of the three data representations—the MA plot, the table, and the heat map—can be further investigated in one of the other representations: A transcript with an interesting differential abundance in the MA plot can be selected to populate the table, a transcript with interesting statistical values in the table can be selected to generate an experiment-wide heat map, and treatments with an interesting transcript abundance in the context of the whole experiment can be selected to generate a new MA plot. Curated lists of transcripts of interest can be generated, added to, and pruned continuously (Figure 1H). The export and import functions (Figure 1I) allow analyses to be continued across datasets and experiments, if transcript names remain consistent across experiments.

### EXAMPLE DATASETS FROM HUMAN, MOUSE, AND WORM

To demonstrate the utility of DrEdGE we have generated three sample DrEdGE websites using data mined from published human, mouse, and worm RNA-seq studies. These examples showcase a range of options for heat map graphics (Figure 2A-E), each designed to suit the different sizes and biological features of the dataset. The final input files for these websites are available in Supplementary File 1, and can be referenced as templates for new DrEdGE websites.

**Figure 2:**
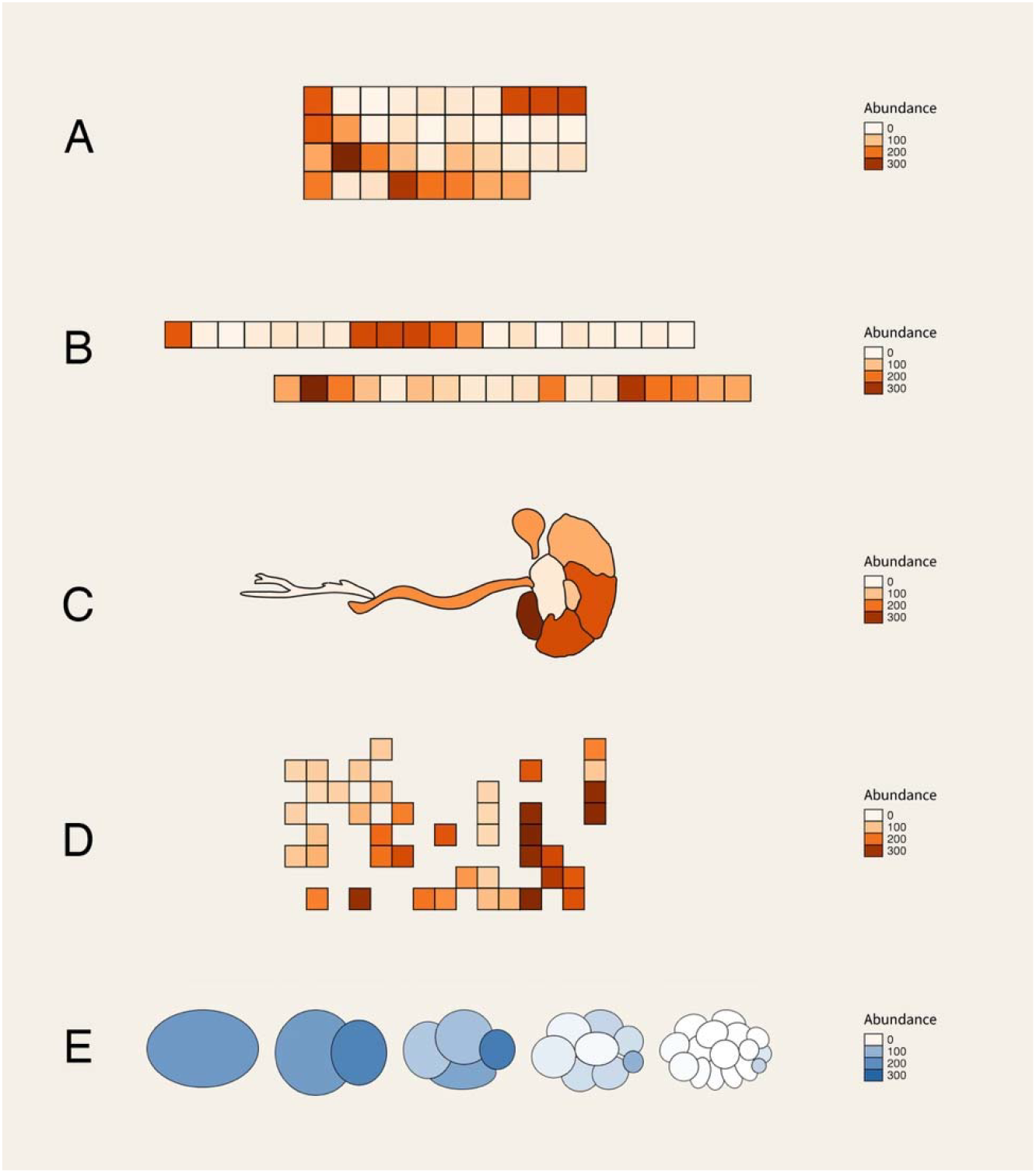
Graphical options for heat map. A) Default box icons in a default compact grid. B) Box icons in a custom linear arrangement. C) Box icons custom-arranged in a grid. D) SVG illustrated icons. E) SVG illustrated icons (from Tintori et al. 2016)^27^.

Although we have generated these examples from publicly available datasets, we strongly encourage the lab generating and analyzing original data to create a DrEdGE website themselves. This will ensure that the website is created with the most thorough understanding of the idiosyncrasies of the data, and the statistics they require.

#### Forty-two whole embryo C. elegans developmental time points

Total RNA-seq data from whole *C. elegans* embryos were mined from Boeck et al, 2016^26^ to create <u>dredge.bio.unc.edu/c-elegans-timecourse</u>. In this study, RNA abundance was reported for forty-two time points from 28 to 737 minutes after the two-cell stage.

Because these samples were numerous and have a linear relationship to each other, we graphically represented them in the heat map using box icons in a custom arrangement, as shown in Figure 2B.

#### Nine neuronal tissues from humans

Human total RNA sequencing data were mined from the ENCODE consortium^4^. We selected nine neuronal tissues—camera-type eye, cerebellum, diencephalon, frontal cortex, occipital lobe, parietal lobe, spinal cord, temporal lobe, and tibial nerve—to generate <u>dredge.bio.unc.edu/human-neuronal-tissue</u>. All libraries were originally prepared and sequenced by the Gingeras lab for the ENCODE consortium.

Because the number of treatments used was relatively small, and because the treatments have a spatial and anatomical relationship to each other, we created custom SVG icons to illustrate each tissue type in the heat map data representation, as shown in Figure 2C.

#### Forty embryonic tissues from mice

Mouse polyadenylated RNA sequencing data were mined from the ENCODE consortium^4^ to create <u>dredge.bio.unc.edu/mouse-embryonic-tissue</u>. Sixteen tissue or cell types were selected: embryonic facial prominence, forebrain, heart, hindbrain, intestine, kidney, limb, liver, midbrain, neural tube, skeletal muscle tissue, spleen, stomach, C2C12 cells, C3H10T1/2 cells, and C3H myocyte cells from C2C12. All samples were prepared and sequenced by the Wold lab for the ENCODE consortium.

Because these samples are numerous we used the default box icons, rather than illustrations, to represent them in the heat map. Because the samples can each be defined by two factors—time point and tissue type—we organized the icons in a custom 2D grid, with each row describing a developmental stage and each column describing a tissue type, as shown in Figure 2D. In some instances, it may be appropriate to show treatments defined in space *and* time with illustrative icons, as in the example from Tintori et al 2016 shown Figure 2E^27^.

### USAGE STEPS FOR GENERATING A DREDGE WEBSITE

Specific detail about the format and function of each file is provided in Supplementary File 2, on the DrEdGE configuration page (http://dredge.bio.unc.edu/blank/?page=configure) and in the instructional video (https://vimeo.com/336521569/816b9585d8). Here we present a brief summary.

#### Coding requirements

We designed the DrEdGE software such that no coding is necessary to create a DrEdGE website, if the author starts with normalized transcript abundance data. If starting from sequence files (.fasta or. fastq), the author must independently align reads to the genome and process the alignments to generate normalized transcript count files.

#### Input files

The DrEdGE software builds a DrEdGE website from five required components, and four optional components (Figure 3, and described in full detail in Supplementary File 2): (1) a table of transcript abundance counts, (2) a table describing the experimental design, (3) a directory of pairwise comparison tables, (4) minimum and maximum values from the comparison tables within that directory, and (5) a JSON file describing the experimental design. The author can provide only the first two components and use R scripts from the DrEdGE package to generate the last three components. Alternatively, if the author has specific statistical methods they prefer to use, the author can generate all five components independently.

**Figure 3:**
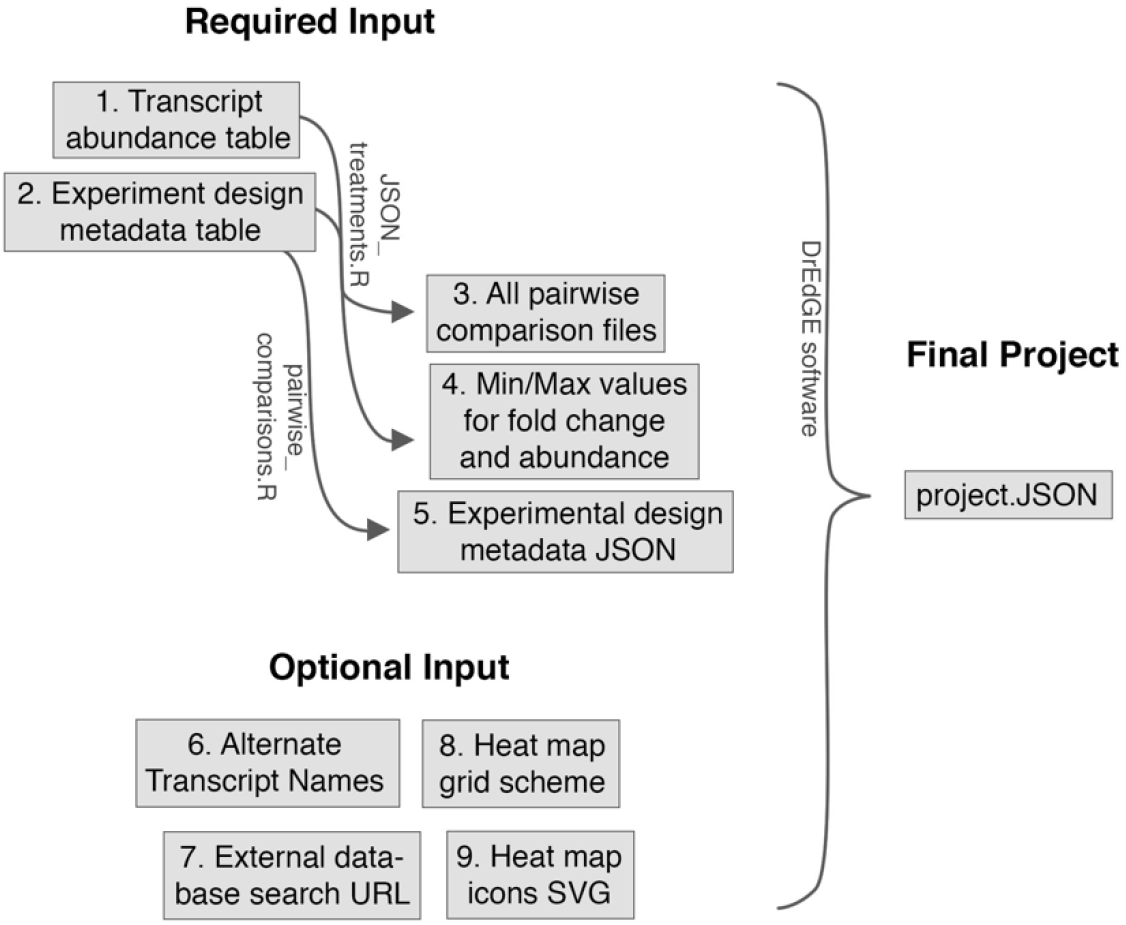
Pipeline of data input and products. Each file in described in further detail in Supplementary File 2.

There are four optional components the DrEdGE software can additionally accept. These include (6) a table of transcript synonyms, or alternate names, (7) a base URL for searching for transcripts on an external organism-specific database, (8) a custom grid scheme for organizing treatments in a 2D heat map grid, and (9) custom SVG icons representing treatments in an illustrated heat map.

#### Processing input files to generate a DrEdGE website

To process the 5-9 components described above into a DrEdGE website, the author must download the DrEdGE package from <u>dredge.bio.unc.edu</u>, unzip the package, and host this DrEdGE directory locally. The index.html file within that directory will present instructions for how to upload each of these components, perform a test run, and save the project once assembly is complete. The author may then upload the directory to the Web to share their DrEdGE website privately with collaborators, or publicly as part of a publication.

#### Storage space required to host DrEdGE website

The final size of the DrEdGE directory will depend primarily on the pairwise comparison tables. If the number of transcripts evaluated (t) is known, and the number of treatments (n) in the experiment is known, the size of the DrEdGE directory can be estimated: Size = (t) * (n^2) * 15 bytes. The three samples presented in this publication range in size from 40 to 500 MB.

## Discussion

As the genomic era ushers in larger and larger datasets, there is a growing need for tools that facilitate data mining and collaboration by sharing genomics datasets in a partially processed, interactive format. Here, we have presented DrEdGE, a program that allows genomicists to easily create interactive data visualization websites. On a DrEdGE website, others can explore a dataset by comparing differential transcript abundance between samples, surveying experiment-wide transcript abundance patterns, and filtering or sorting by specific statistics. This tool allows genomics researchers to share their work with other interested parties, either during collaboration or for data mining after publication.

For simplicity, we have limited our demonstration of DrEdGE’s utility to RNA-seq experiments. DrEdGE’s flexible design, however, allows users to visualize any two-dimensional numeric dataset, including comparative proteomics, population diversity maps, or even non-biological datasets.

### Iterative analysis

DrEdGE’s iterative analysis design allows readers to select data points of interest from each data representation to investigate further in the next representation. The reader can curate a list of transcripts through multiple analyses. This allows the reader to ask sophisticated biological questions of the data, and generate statistically supported results. This differs from current interactive genomics data visualization websites, such as genome browsers, because comparative biological questions can be asked, tested, refined, and tested again continuously. Exporting curated lists of transcripts from one experiment and importing it to another experiment also allows for integrated analyses across different datasets or types of assays.

### Statistical flexibility and low technical overhead

The potential for authors to use statistical methods of their choice has two benefits. First, it means that DrEdGE consists solely of static files, making it easier to host and maintain than a typical dynamic Web application powered by a database and server-side program (written in e.g. PHP, Python, Perl)^29^. Second, it allows DrEdGE to fit within diverse data analysis pipelines. Each experiment requires unique statistical considerations, whether due to idiosyncrasies of the genome, the molecular technique, or the philosophy of the researcher, which DrEdGE is able to accommodate. This flexibility expands DrEdGE’s utility in a rapidly changing field.

## Conclusions

We have presented DrEdGE, a tool for generating interactive visualizations of large genomics datasets. These visualizations allow interested parties to independently explore data by generating their own MA plots, transcript abundance statistics tables, and experiment-wide heat maps, and to modify lists of transcripts of interest through multiple analyses. This tool fills the gap in data sharing practices between raw data on databases (that require a substantial amount of time and expertise to process) and a handful of completed analyses in publications (that are static and cannot be queried further). By removing the technical obstacles that prevent interested parties from exploring multivalent datasets, DrEdGE can increase the utility and impact of these datasets.

## Methods

### Implementation and availability

DrEdGE is a browser-based application powered by client-side JavaScript. The entire application is bundled into one HTML file and one JavaScript file, using the JavaScript compiler Browserify^30^ and the automation tool GNU Make^31^. All of the pages in the application are built and rendered using the React library^32^. Interactive data visualizations are created using the D3.js library^33^. All code is freely available on GitHub (<u>github.com/ptgolden/dredge</u>) under the GNU Affero General Public License (AGPLv3).

### Sourcing of sample datasets

RNA-seq data for humans, mice, and worms were collected from ENCODE^4^ or Boeck et al., 2016^26^. Human experiment accession numbers: ENCSR000AFO, ENCSR000AEW, ENCSR000AEX, ENCSR000AEY, ENCSR000AFD, ENCSR000AFE, ENCSR000AFJ, ENCSR858QEL, ENCSR648OSR, ENCSR796HLX, ENCSR272UNO, ENCSR000AFH. Mouse experiment accession numbers: ENCSR809VYL, ENCSR851HEC, ENCSR823VEE, ENCSR636CWO, ENCSR160IIN, ENCSR970EWM, ENCSR752RGN, ENCSR691OPQ, ENCSR284YKY, ENCSR727FHP, ENCSR597UZW, ENCSR526SEX, ENCSR420QTO, ENCSR848GST, ENCSR537GNQ, ENCSR173PJN, ENCSR347SQR, ENCSR830IVQ, ENCSR648YEP, ENCSR448MXQ, ENCSR867YNV, ENCSR826HIQ, ENCSR096STK, ENCSR457RRW, ENCSR992WBR, ENCSR307BCA, ENCSR908JWT, ENCSR343YLB, ENCSR557RMA, ENCSR719NAJ, ENCSR337FYI, ENCSR115TWD, ENCSR667TOX, ENCSR946HWC, ENCSR579FCW, ENCSR290RRR, ENCSR178GUS, ENCSR000AHY, ENCSR000AHZ, ENCSR000AHX, ENCSR000AIA. Counts were compiled into transcript count abundance tables and experimental design tables in R. Transcript synonym tables for humans, mice, and worms were generated from the HUGO Gene Nomenclature Committee’s Biomart^34,35^, Mouse Genome Informatics^36^, and Wormmine^37^, respectively.

## DECLARATIONS

### Availability of data and material

The datasets used in this study are all previously published (Boeck et al. 2016, The ENCODE Project Consortium 2012, and Tintori et al. 2016). The reformatted and processed data used to create the three example DrEdGE websites are available in Supplementary File 1. Original code for the DrEdGE program is available at dredge.bio.unc.edu and github.com/ptgolden/dredge.

### Authors’ contributions

Project was conceived of by ST, and designed by PG and ST with input from BG. Javascript code by PG, R code by ST. Manuscript written by ST with input from PG and BG.

## Acknowledgements

The authors would like to thank Dan McKay, Jonathan Seguin, Max Boeck, David King, Ann Wehman, and Kira Heikes for their thoughtful feedback on the preprint of this manuscript.

## SUPPLEMENT

**Supplementary File 1: Materials used to generate the three example DrEdGE websites** Because these data files cannot be uploaded at Word or LaTex files, we are providing an external link to the source data tables. It is a 240 MB zipped file. https://www.dropbox.com/s/la89lezajk2c1pp/Supplement%201%20-6%20source%20files.zip?dl=0

**Supplementary File 2: Descriptions of input files for generating a DrEdGE website**

All files and numbers correspond to Figure 3.

### 1 Transcript abundance table

The table of normalized transcript counts must have one row for each transcript, and one column for each individual replicate in the experiment. Row names (transcript IDs) must be unique, and column names (replicate IDs) must be unique and match the experimental design metadata (Files #2 and #5).

### 2 Experimental design metadata table

The experimental design table must include a row for each replicate, and must include the following columns: “replicate.id”, “treatment.id”, and “treatment.name”. Replicate IDs must match the column names of the transcript abundance table (File #1). Treatment names should be meaningful to a new reader exploring the DrEdGE website. No row names are necessary.

### 3 Directory of all pairwise comparison data tables

This directory must contain a unique file for each unique pair of treatments, including self-to-self comparisons. Each file must be a table with a row for each transcript, and the following three columns in this order: (1) the logarithm of the fold change in transcript abundance between the first and second treatment, (2) the logarithm of the average transcript abundance amongst all replicates of both treatments, and (3) the P-value for fold change described in column 1. Row names (transcript IDs) must match the row names in the transcript abundance file (File #1). The names of the files must reference the treatment IDs used in the experimental design metadata (Files #2 and #5). The author is encouraged to use his or her own preferred analytic methods to create the pairwise comparison tables, but is also welcome to run the pairwise_comparisons.R script provided in the DrEdGE package. This script receives the transcript abundance table (File #1) and the experimental design file (File #2) as input, and uses edgeR to generate, name, and export all possible pairwise comparisons. The edgeR analysis models the data as a negative binomial distribution, and models overdispersion as a Poisson model. An empirical Bayes procedure is used to moderate the degree of overdispersion across transcripts, and differential expression is assessed using an exact test adapted for overdispersed data^28^.

### 4 Minima and maxima from pairwise comparison tables

These minima and maxima will be used to set the default axis limits for the MA plots. The numbers will be entered manually, so format is not important. If the author has run the pairwise_comparisons.R script to generate the directory of pairwise comparison tables (Files #3), these minima and maxima values will automatically be calculated and a report will be generated.

### 5 Experimental design metadata JSON

The experimental design metadata (File #2) must also be provided as a JSON (JavasScript Object Notation) file in which each first-order object describes each treatment from the experiment. The JSON_treatments.R script provided in the DrEdGE package will accept File #2 and generate this JSON file. If the author wishes to generate the JSON file independently, the first order object keys must be the short unique codes for the treatment (“treatment.id” in File #2), and the values must include the following second order objects: (1) A "label" key that points to a verbose name for the treatment (“treatment.name” in File #2), and a "replicates" key that points to the full list of replicate IDs for the treatment (“replicate.id” in File #2).

### 6 Transcript synonyms or alternate names

The author may wish to associate a preferred “readable” name with each transcript, and include other code names that readers might use in the search bar. In this case the author may include a file of alternative names, or synonyms, for transcripts. With one line of the file per transcript, each line must begin with the transcript name that the DrEdGE visualization will show. Following that first-choice name should be a list all other names or codes for the transcript, separated by tabs. In the input files for our three example websites (Supplementary File 1), we have included transcript synonym tables for human, mouse, and worm.

### 7 Organism-specific database search URL

If the author would like transcripts to link to an external database (i.e. Wormbase^37^), he or she may provide a generic search URL for this database, with “%name” inserted where the transcript name would be (e.g. http://wormbase.org/search/gene/%name).

### 8 Custom grid layout for grid heat map

The default settings represent each treatment as a box icon, with boxes fit together in a compact grid (Figure 2A). Alternatively, the author can specify how he or she would like these boxes arranged (Figure 2B,D). To do this, the author must create a comma delineated file in which each row of text describes the sequence of treatments (or white space) for each row of the grid.

### 9 Custom SVG for illustrated heat map

The author can display the heatmap as a graphic illustration by uploading an SVG (Scalable Vector Graphics) file with custom icons representing each sample (Figure 2C,E). The author must create a vector file with one object for each treatment, and each object must be given a name that matches the treatment ID used in the experimental design metadata files (Files #2 and #5).

